# High quality whole genome sequence of an abundant Holarctic odontocete, the harbour porpoise (*Phocoena phocoena*)

**DOI:** 10.1101/246173

**Authors:** Marijke Autenrieth, Stefanie Hartmann, Ljerka Lah, Anna Roos, Alice B. Dennis, Ralph Tiedemann

**Author notes:** Corresponding author: Prof. Dr. Ralph Tiedemann.

## Abstract

The harbour porpoise (*Phocoena phocoena*) is a highly mobile cetacean found in waters across the Northern hemisphere. It occurs in coastal water and inhabits water basins that vary broadly in salinity, temperature, and food availability. These diverse habitats could drive differentiation among populations. Here we report the first harbour porpoise genome, assembled *de novo* from a Swedish Kattegat individual. The genome is one of the most complete cetacean genomes currently available, with a total size of 2.7 Gb and 50% of the total length found in just 34 scaffolds. Using the largest 122 scaffolds, we were able to validate a high level of homology to the chromosome-level genome assembly of the closest related species for which such resource was available, the domestic cattle (*Bos taurus*). The draft annotation comprises 22,154 predicted gene models, which we further annotated through matches to the NCBI nucleotide database, GO categorization, and motif prediction. To infer the adaptive abilities of this species, as well as their population history, we performed a Bayesian skyline analysis, and produced results that are concordant with the demographic history of this species, including expansion and fragmentation events. Overall, this genome assembly, together with the draft annotation, represents a crucial addition to the limited genetic markers currently available for the study of porpoises and Phocoenidae conservation, phylogeny, and evolution.

## Introduction

A central question in evolutionary biology is understanding the genomic basis of key innovations and adaptations, the respective impacts of selection *vs*. drift on specific genes, and how these patterns vary across the entire genome. In non-model organisms, these questions have been typically addressed using just a few genetic markers (e.g. microsatellites, single mitochondrial or nuclear genes), often in the absence of knowledge about the genomic position of the analyzed loci (e.g., Diefenbach et al., 2015; Ellegren & Galtier, 2016; Hoelzel et al., 1999; Lopes et al., 2015; Wiemann et al., 2010). However, in the last ~10 years the cost of sequencing has decreased, while read quality and sequence length have increased (Neely et al., 2018), and genome-wide resources are no longer limited to a few model organisms. With these resources, it is now possible to detect more subtle differences among species and populations and to extend these beyond nucleotide polymorphisms to larger structural variants, linkage groups, and highly divergent genes and regions on a genomic scale (e.g. Johannesson et al., 2017; L. F. Li et al., 2017; Whiteman, 2017; Willoughby et al., 2017).

Among the non-model organisms, cetaceans are a valuable group of animals to invest in creating solid genetic resources; cetaceans attract considerable attention, both scientifically and in the general public, and genomic resources enable further examination of their adaptive evolution and facilitate population assessments in the context of conservation and management. As fully aquatic mammals, cetaceans encounter escalating levels of mostly human-caused threats that lead to death, infertility, and reduction of population size and stability (Fietz et al., 2013; Godard-Codding et al., 2011). These include noise pollution (Dyndo et al., 2015; Nabe-Nielsen et al., 2014), marine debris and by-catch (Scheidat et al., 2008; Unger et al., 2017), and infectious diseases (Siebert et al., 2001; van Beurden et al., 2017).

Beside acoustic (Amundin, 2016) and photo-id monitoring, a number of genetic markers are frequently and intensively used to estimate population status and boundaries. The most commonly implemented markers are microsatellites (e.g., Fontaine et al., 2010; Hoffman & Amos, 2005; Palo et al., 2001) and mitochondrial DNA (e.g., Autenrieth et al., 2018; Graves et al., 2009; Speller et al., 2016), and more recently genome-wide single nucleotide polymorphism (SNP) surveys (e.g., Lah et al., 2016). New genomic resources hence do not only facilitate evolutionary research, but enable elevation of conservation genetics approaches to the genomic level (Foote et al., 2015).

Genetic resources available for cetaceans are growing, but do not yet adequately cover the diversity of this group (Foote et al., 2016; Keane et al., 2015; Nery et al., 2013; Sun et al., 2013; Yim et al., 2013; Zhou et al., 2013). Thus far sequenced genomes include three baleen whales (DeWoody et al., 2017; Kishida et al., 2015; Yim et al., 2013) and five species (with a collective seven genomes) of the toothed whales (Foote et al., 2015; Jones et al., 2017; Lindblad-Toh et al., 2011; Neely et al., 2018; Warren et al., 2017; Zhou et al., 2013), but the completeness of these assemblies varies (Table 1). Within the odontoceti only four of the ten families (*Physeteroidae*, *Lipotidae, Monodontidae* and *Delphinidae*) have been whole-genome sequenced to date. The family *Phocoenidae*, sister group to the *Monodontidae*, encompasses seven species, three of them threatened (Gatesy et al., 2013; Geisler et al., 2011, Rojas-Bracho et al. 2017, Wang et al., 2017). Being the smallest cetaceans in stature, phocoenids occur close to the coast and are therefore, like river dolphins, especially prone to human induced threats. This puts these species particularly at threat of decline or even extinction, as in the case of the Vaquita, endemic to Mexico (Taylor et al., 2017); yet no full genomic resource is available for these species.

**Table 1.** Publicly available cetacean genomes in comparison to the new genome of the harbour porpoise (*Phocoena phocoena*)

| Species name | NCBI-ID | WGS project | total sequence length in Gb | # of scaffolds | scaffolds N50 | Genome coverage | publication |
| --- | --- | --- | --- | --- | --- | --- | --- |
| *Phocoena phocoena* | Pho_pho_vs1_UP | tba | 2.68 | 13,498 | 23,753,446 | 87.0x | this study |
| *Delphinapterus leucas* | ASM228892v2 | NQVZ01 | 2.36 | 6,972 | 19,885,328 | 117.0x | Jones et al. (2017) |
| *Tursiops truncatus* | NIST_Tur_tru v1 | MRVK01 | 2.13 | 2,648 | 26,555,543 | 114.5x | Neely et al. (2018) |
| *Tursiops truncatus* | Ttru_1.4 | ABRN02 | 2.55 | 240,558 | 116,287 | 30x | Lindblad-Toh et al. (2011) |
| *Orcinus orca* | Oorc_1.1 | ANOL02 | 2.37 | 1,668 | 12,735,091 | 200.0x | Foote et al. (2015) |
| *Lipotes vexillifer* | Lipotes_vexillifer_v1 | AUPI01 | 2.43 | 30,713 | 2,419,148 | 115x | Zhou et al. (2013) |
| *Physeter macrocephalus* | Physeter_macrocephalus-2.0.2 | AWZP01 | 2.28 | 11,711 | 427,290 | 75x | Warren et al (2017) |
| *Physeter macrocephalus* | ASM283717v1 | PGGR01 | 2.52 | 59,237 | 37,183,148 | 248x | ds (Fan et al. 2017) |
| *Eschrichtius robustus* | ASM218922v1 | NIPP01 | 2.85 | 57,203 | 187,455 | 11.0x | DeWoody et al. (2017) |
| *Balaenoptera acutorostrata* | BalAcu1.0 | ATDI01 | 2.43 | 10,776 | 12,843,668 | 92x | Yim et al (2014) |
| *Balaenoptera bonaerensis* | ASM97880v1 | BAUQ01 | 2.23 | 421,444 | 20,082 | 60x | Kishida et al. (2015) |

NCBI-ID: assembly name, WGS project: whole genome sequence project, publications: ds = direct submission, not yet linked to a publication.

**Table 2.** Summary statistics of sequence data used for the draft and final HiRise assemblies.

|  | Insert (bp) | n reads | Coverage |
| --- | --- | --- | --- |
| Illumina Library 300 | 300 | 794 M | 74.1 X |
| Illumina Library 500 | 500 | 473 M | 44.2 X |
| Chicago library | 1-50kb | 528 M | 87.2 X |

We have chosen to develop a high quality genetic resource of the harbour porpoise (*Phocoena phocoena*). This cetacean species occurs from sub-polar to temperate coastal waters in the Northern hemisphere (Fontaine et al., 2017; Gaskin, 1984; Lah et al., 2016). As an apex predator, the harbour porpoise (*Phocoena phocoena*) is a key indicator for conservation and biodiversity measurements in the Nordic Seas (Hooker & Gerber, 2004; Lawrence et al., 2016; Sergio et al., 2008).

The population size of harbour porpoises in European Atlantic Shelf waters is estimated to be 375,000 with shifts across the last decade in the exact regions they occupy (e.g. in the North Sea; (Hammond et al., 2013)). Estimates of population size in the western Baltic Sea are smaller at approximately 40,000 animals (Benke et al., 2014; Scheidat et al., 2008; Viquerat et al., 2014), while the Baltic Sea proper only has an estimated < 500 individuals (Amundin, 2016), classifying this population as critically endangered (Benke et al., 2014; Hammond et al., 2008; Scheidat et al., 2008).

In addition to occupying different habitats, these populations differ in a number of features, including food preference (Hammond et al., 2013), activity patterns (Nuuttila et al., 2017), and fine scale morphological features. These differences are corroborated by genetic differentiation (Fontaine et al., 2012, 2014; Wiemann et al., 2010) and significant isolation by distance (Lah et al., 2016). This led to the suggestion that different ecotypes of harbour porpoise in the North Atlantic and Baltic Sea exist (Fontaine et al., 2014, 2017; Galatius et al., 2012). Given this evidence for genetic differentiation among populations, a genomic resource would not only be a valuable addition to the few existing cetacean genomes, but it would also be very useful for monitoring the separate porpoise populations and investigating whether there are specific local adaptations, e.g., to differences in temperature and salinity, in these small cetaceans.

We present here the first *de novo* assembly of the full genome of the harbor porpoise, scaffolded with *in vitro* proximity ligation data (hereafter “Chicago” library), and draft-annotated to predict its coding proteins and their functions (Deposited at NCBI as BioProject: PRJNA417595 with BioSample-ID: SAMN08000480). We demonstrate a high level of completeness of the assembly, showing that many of our scaffolds are near-chromosome-level, and provide insight into past demographic dynamics of the population the sequenced individual originated from, using a Bayesian skyline plot (Li & Durbin, 2011).

## Materials and Methods

### DNA sampling

Tissue for whole genome sequencing came from a single individual from the Kattegat sea (Glommen - Falkenberg), Sweden (ID: C2009/02665). Muscle tissue was sampled in July 2009 from a by-caught female of probably young age (22.4kg, 110.5m), frozen and transported to Potsdam, Germany for DNA extraction. Sample preparation and Genomic DNA isolation were performed following the Quiagen DNeasy Blood & Tissue Kit (Cat 69506, Hilden). Successful high molecular weight DNA-isolation was confirmed by Sanger sequencing of the mitochondrial control region and visualization of fragment sizes of the entire extraction using a Tape Station (Agilent 2200, Santa Clara, CA 95051). By mtDNA control region sequencing we verified that the analyzed specimen carried haplotype PHO7 (Tiedemann et al., 1996), indicative of the separate Beltsea population of the Kattegat/Western Baltic Sea region (Lah et al., 2016; Wiemann et al., 2010).

### Genome sequencing and assembly

The draft *de novo* assembly was constructed from two libraries (insert sizes ca. 300 and ca. 500bp); sequenced in 125bp PE reads on the Illumina HiSeq 2500 at EUROFINS Genomics. Between 90.43% and 91.52% of the bases had a quality of 30 or higher; the mean quality values ranged between 34.39 and 34.78. Reads were trimmed using Cutadapt v1.10 (Martin, 2011), and an initial assembly was performed using SOAPdenovo2 (Luo et al., 2015). DNA from the same sample was used by Dovetail Genomics for construction of the Chicago library (Putnam et al., 2016) and sequenced in 150bp PE reads on an Illumina NextSeq500 at the University of Potsdam. The draft assembly was then scaffolded with the Chicago library data using the HiRise pipeline (Dovetail Genomics LLC, Santa Cruz, CA, USA).

Blobtools was run to examine potential contaminants, based on divergence in GC-content and read coverage variation across the assembly (Laetsch & Blaxter, 2017). Presence of core, single copy, and orthologous genes was measured using CEGMA and BUSCO with the latter run in the genome mode for the Laurasiatheria database (Simão et al., 2015).

### Genome annotation

Genome annotation was performed using Maker2 (Holt & Yandell, 2011), which makes use of different programs and draws from several lines of evidence. Prior to annotation, repetitive elements were soft-masked with RepeatMasker (Smit et al., 2013-2015) using the te_protein repeat database (Smith et al., 2007). In the first Maker2 run, three gene predictors were used: SNAP (Bromberg et al., 2008) was *ab initio* trained with the CEGMA results (Parra et al., 2007), Genemark-ES (Ter-Hovhannisyan et al., 2008) was run using an Hmm produced by *ab initio* training on the whole *P. phocoena* genome, Augustus was run using the presets for human, as is recommended for vertebrates (Stanke et al., 2004). Protein sequences, supplied as evidence, were obtained from the complete UniProt database (553,941 Proteins) plus NCBI entries of 184,527 proteins (retrieved on 20 March 2017) with matches to the following keywords: “*Balaenopteridae*”, “*Lipotes vexillifer*”, “*Neophocaena*”, “*Orcinus orca*”, “*Phocoena*”, “*Physeter catodon*”, “*Pontoporia blainvillei*”, “*Tursiops truncatus*”.

For the second Maker2 run, we created a new SNAP-Hmm based on the first Maker2 output and ran it with the same parameters as the first run, exchanging only the SNAP Hmm and excluding the protein evidence. The resulting CDS predictions were extracted from the final gff file, which was created by *fathom* implemented in SNAP (Bromberg et al., 2008). These gene predictions were further verified by a Blastn search against the entire GenBank non-redundant (nr) nucleotide sequence database (date downloaded 21.07.2017) using a maximum number of 3 target sequences and an e-value of 1e-10. Summary statistics were generated using Genome Annotation Generator (Hall et al., 2014). We then used all CDS and their Blast results in Blast2go (Goetz et al., 2008) to identify conserved protein domains with InterProScan. We functionally annotated the CDS with GO terms, which are a controlled vocabulary to further categorize gene function (Ashburner et al., 2000; Carbon et al., 2017).

### Comparative genomics

The closest relative to *P. phocoena* with a chromosome-level assembly currently available is the domestic cattle, *Bos taurus*. To validate our assembly, we compared our scaffolds to the *B. taurus* chromosomes (assembly UMD 3.1.1 downloaded from NCBI, ACCESSION DAAA00000000). Specifically, the 122 *P. phocoena* scaffolds of at least 1Mbp were aligned to the *B. taurus* chromosomes using the nucmer software of the MUMmer package v. 3.23 (Kurtz et al., 2004). From the coordinates of these alignments, runs of ten or more consecutive matches of each at least 250bp between a given *P. phocoena* scaffold and a *B. taurus* chromosome were extracted using custom perl scripts. Their start and end positions were used to generate a Circos plot (http://circos.ca/) that shows regions of collinearity as well as rearrangements. For the Circos plot, separate ribbons are displayed between a *B. taurus* chromosome and a *P. phocoena* scaffold for consecutive hits that were each no more than 20,000 bp apart. If a hit was more than 20,000bp from the next run of consecutive hits, a new ribbon was started (Figure 1).

**Figure 1.**
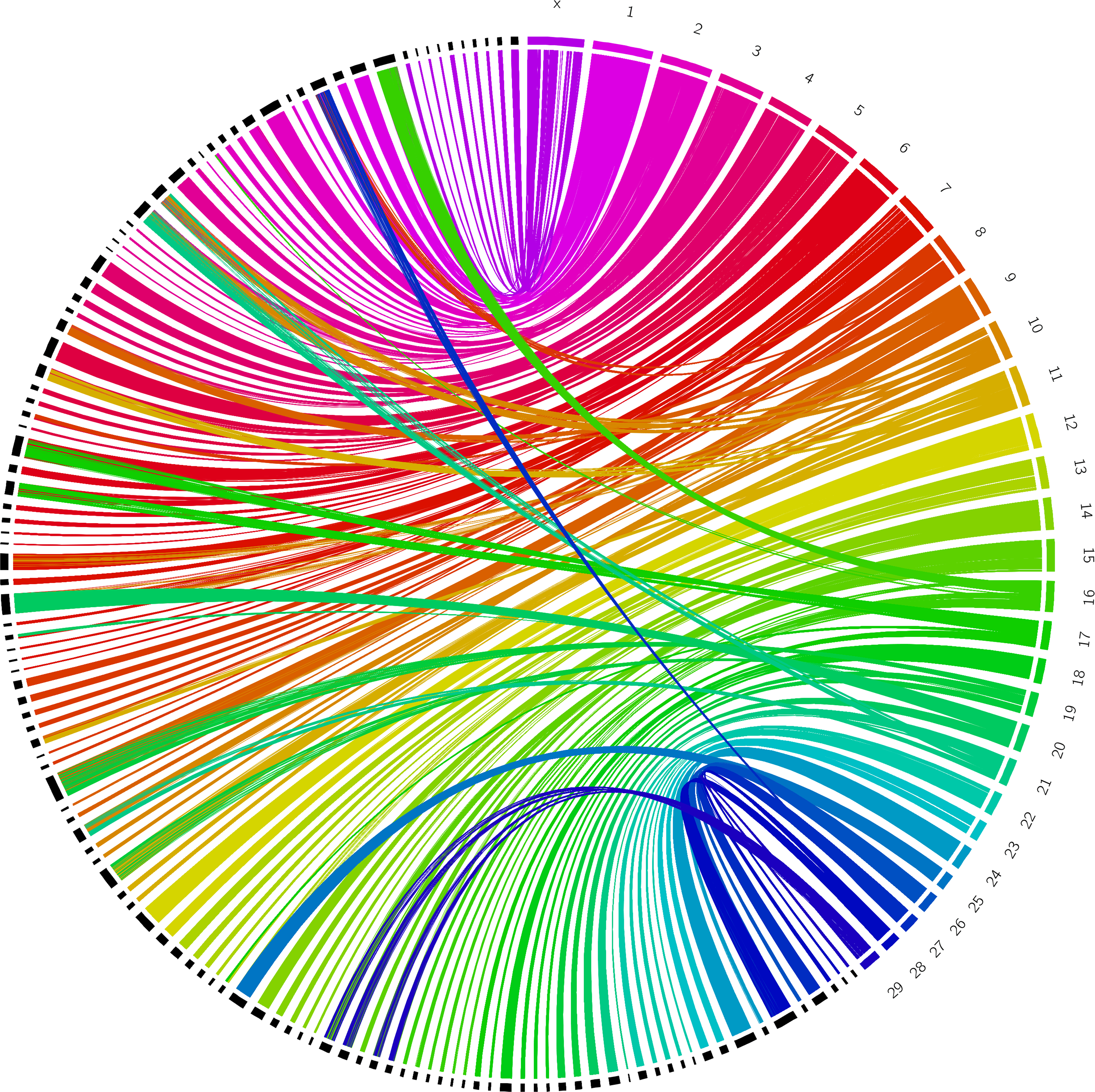
Comparison *of P. phocoena* scaffolds and *B. taurus* chromosomes. Each *B. taurus* nuclear chromosome is shown in a different colour on the right outer rim; they are labelled X for the X chromosome and 1-29 for the 29 autosomes. *P. phocoena* scaffolds of at least 1 Mbp are shown in black on the left outer rim and not labeled. Only the 122 largest scaffolds with consecutive MUMmer hits (as described in the main text) are included. Matches between *B. taurus* chromosomes and *P. phocoena* scaffolds are shown in the colour of the *B. taurus* chromosomes.

### Demographic history

Using genome-wide diploid sequence data, it is possible to reconstruct the population history by estimating population sizes in the past (Li et al., 2011). To estimate the demographic history of the sequenced individual, phased sequence data is required. Since we did not have this, we generated the diploid sequence data as follows: we used the SNP frequency spectrum based on our genome assembly, which is a haploid sequence, and the PE reads used to construct the *de novo* assembly prior to Chicago scaffolding (described above, we used both insert sizes). These reads were first mapped back to the final assembly using BWA (Li & Durbin, 2009). SNP data was extracted from the resulting bam files, and variants were extracted using samtools vs.1.6. (Li, Handsaker, et al., 2009) and bcftools (Li, Handsaker, et al., 2009). Results were then used to generate the final Bayesian skyline plot in the PSMC package, using perl scripts psmc2history.pl and psmc_plot.pl (Li et al., 2011) and including 100 bootstraps (Li et al., 2011). The parameters of the PSMC analysis were set following the recommendation from the authors (Li & Durbin, 2011, https://github.com/lh3/psmc) and we assumed a generation time of 10 years (Birkun Jr. & Frantzis, 2008) and a mutation rate of 2.2 × 10^−9^ year/site (Taylor et al., 2007).

## Results

### De novo assembly of the P. phocoena genome

Shotgun sequencing produced a total of 1,268M reads (Table 1); these were used to generate a draft assembly with 2.4M sequences (including contigs and scaffolds) and an N50 of 33.1kb. This assembly was combined with the Chicago library data (528M read) for final scaffolding by Dovetail Genomics (Putnam et al., 2016). The final HiRise assembly contains 2,025,248 scaffolds in total and including singletons, with 13,498 scaffolds bigger than 1kb (Table 3), had a total length of 2.7Gb (N50 of 23.8Mb), and was covered by 87x sequence data. The greatest improvements from the addition of the Chicago libraries was in building up the 34 longest scaffolds, which make up approximately half of the entire assembly (Table 3). The BUSCO and CEGMA analyses also suggest that we have largely reconstructed the entire genome, as we identified 96.9% (91.3% complete) of the 2,586 Eukaryotic and 94.2 (88.7% complete) of the 6,253 Laurasiatheria BUSCO core genes and 90% of the 248 ultra-conserved CEGMA genes (54% complete).

**Table 3.**
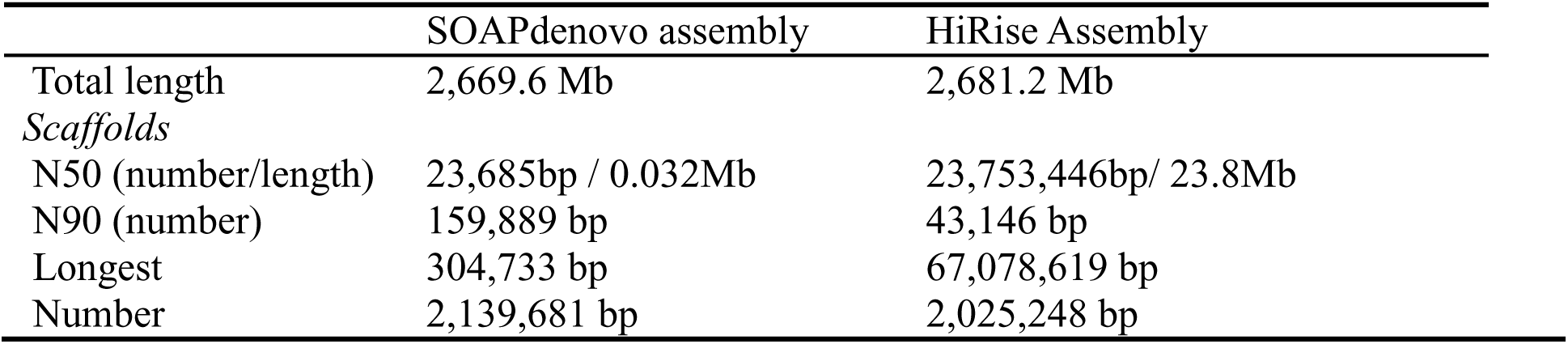
Assembly statistics of the harbour porpoise genome assembly.

|  | SOAPdenovo assembly | HiRise Assembly |
| --- | --- | --- |
| Total length | 2,669.6 Mb | 2,681.2 Mb |
| *Scaffolds* |  |  |
| N50 (number/length) | 23,685bp / 0.032Mb | 23,753,446bp/ 23.8Mb |
| N90 (number) | 159,889 bp | 43,146 bp |
| Longest | 304,733 bp | 67,078,619 bp |
| Number | 2,139,681 bp | 2,025,248 bp |

### Genome completeness and annotation

The Maker2 annotation resulted in a predicted 22,154 coding genes (Table 4). In total, 21,750 CDS had a Blast hit against the nucleotide database, which accounts for 98% of the total CDSs. Of these Blast hits, 99% were from vertebrates, and these were dominated (90%) by hits to cetacea (thereof 59% *Tursiops truncatus*, 27% *Orcinus orca*). Further annotation with InterProScan revealed 250,126 features of these predicted proteins. These comprise hits in several protein domain databases, e.g. 23,319 PFAM protein domains, 37,046 PANTHER gene families, 24,538 SUPERFAMILY annotations and 31,114 Gene3D domains. Assignment of the Blast results to Gene Ontology (GO) categories resulted in 55,143 hits across the GO categories (Figure 2).

**Figure 2.**
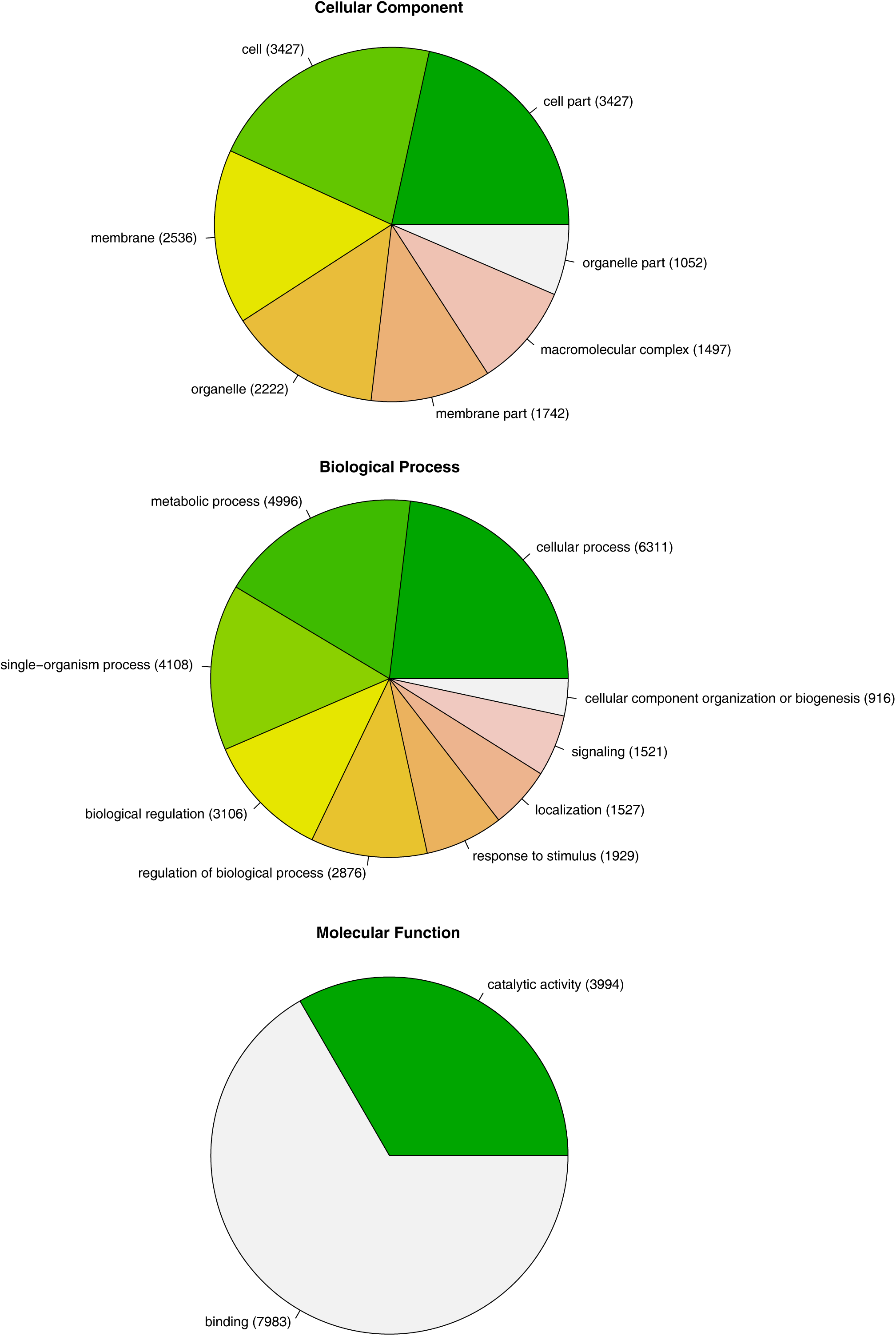
GO-Terms; GO annotation level 2, separated by gene ontology terms: “Cellular Component”, “Biological Process”, and “Molecular Function”. Separate categories are listed, with the number of hits in parentheses.

**Table 4.** Genome annotation statistics

|  | Exons | Introns | Genes | CDS |
| --- | --- | --- | --- | --- |
| Number | 171,735 | 149,581 | 22,154 | 22,154 |
| Longest (in bp) | 10 kb | 1,532 kb | 2,299 kb | 21 kb |
| Mean length | 166 | 3,966 | 28,051 | 1,282 |
| % genome covered by | - | - | 23.2% | 1.1% |
| GC% in CDS | - | - | - | 54.48% |

### Comparison to a chromosome-level assembly

To illustrate the completeness of the harbour porpoise genome, we mapped the 122 biggest scaffolds against the domestic cattle chromosomes, the closest available genome with chromosomal information. Between the 30 domestic cattle chromosomes and the 122 harbour porpoise scaffolds, a total of 24,394 separate matches (as described in the Materials and Methods section) were constructed. The matching regions show some rearrangement (Figure 1). In some cases, ribbons of different colors (coming from different domestic cattle chromosomes) match a single porpoise scaffold. For example, one porpoise scaffold (Sc_11228) encompasses parts of the cow chromosome 1 (violet), cow chromosome 8 (red) and cow chromosome number 27 (blue, Figure 1). The scaffold IDs that match each domestic cattle chromosome are listed in Supplemental Material 1. We find almost complete matching of cow chromosome to a single porpoise scaffold for cow chromosome 25/ porpoise scaffold Sc_12150 (cyan-ribbon). Additionally cow chromosomes 12, 23, 24, 26 and 27 are each matched by only two porpoise scaffolds. Other chromosomes are matched by three or more scaffolds, with a maximum of ten (x-chromosome; Supplement Material 1).

### Inference of Kattegat/Baltic population history

We inferred the demographic history of the harbour porpoise *P. phocoena* Kattegat/Baltic population based on one single individual (Li et al., 2011) using the PSMC algorithm and all of our PE read data. In the resulting graph (Figure 3), the inferred population size (*N_e_*) began to increase around 1.5 Myr ago, reaching an *N_e_* of 500,000 during the following 1 Myr. The estimated population size peaked approximately 200,000 years ago at an estimated *N_e_* ~500,000, before it dropped to a fifth of the original size around 100,000 years ago, leading to a lower *N_e_* of 100,000.

**Figure 3.**
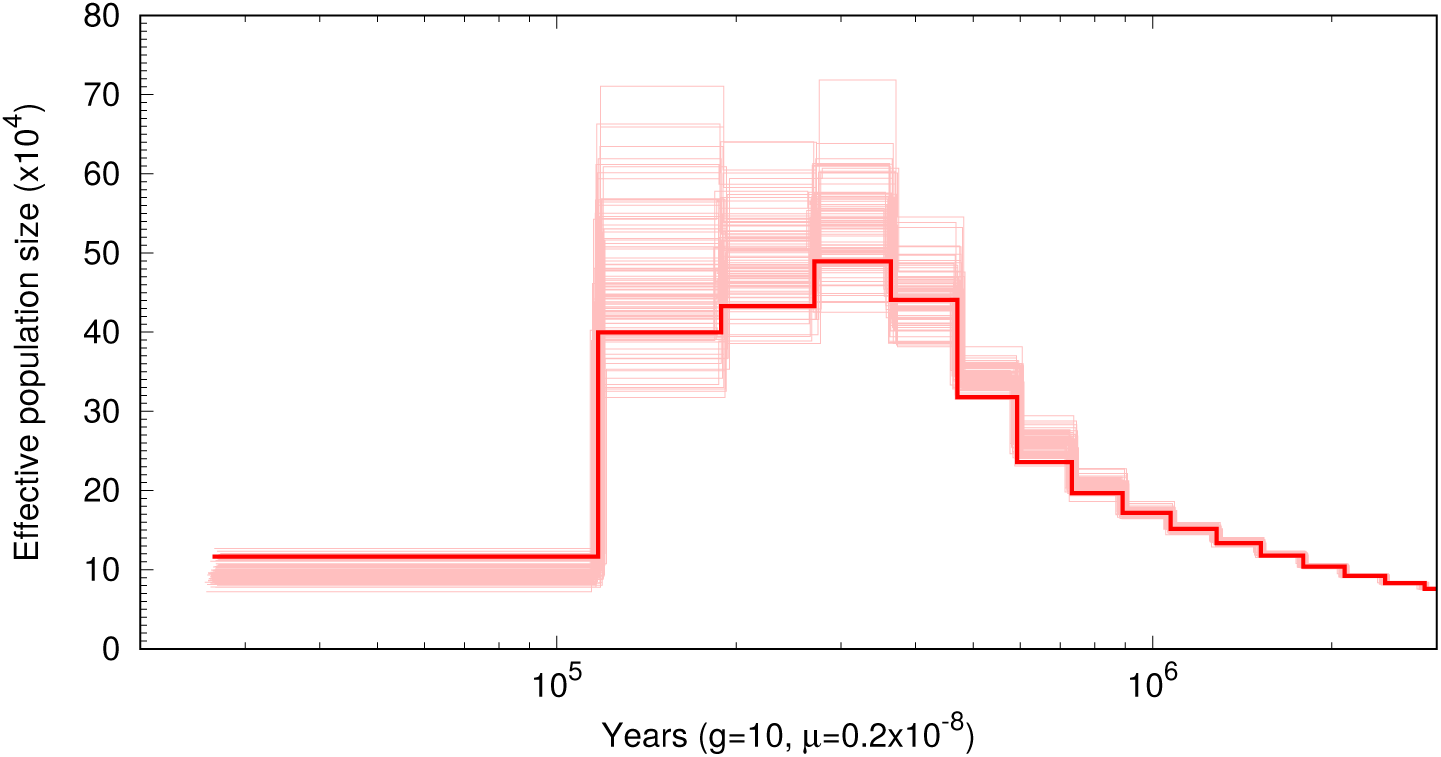
PSMC estimated harbour porpoise population size changes over time for the Baltic Sea. g = generation time; μ = mutation rate (per site, per year). Porpoise data generated on the basis of mapping PE reads to whole genome scaffolds during SNP calling. Pale lines show the trajectories of 100 bootstrap replicates.

## Discussion

We present here a high quality *de novo* genome assembly for the harbour porpoise *Phocoena phocoena*. Scaffolding with Chicago Library method (i.e. *in vitro* proximity ligation) has previously produced high quality genomes in plants (Moll et al., 2017; Paajanen et al., 2017), insects (Nosil et al., 2018; Wachi et al., 2017), and birds (Holt et al., 2018). We have applied this method to a sample from a deceased by-caught cetacean specimen, demonstrating that this approach can be successfully used for non-model mammal species and samples of lower quality, as is typical for cetacean research samples that are often obtained from stranded or by-caught individuals (e.g., Autenrieth et al., 2018; Ottewell et al., 2016; Sabatier et al., 2015; Squadrone et al., 2015).

With a GC-content of 41.4% and a total length of 2.7 GB, this assembly is comparable to other high quality cetacean genomes (Table 1; Groenen et al., 2013; Zimin et al., 2009). In support of this, the scaffold N50 and sequence coverage are similar to the recently published high quality genomes of the beluga whale (Jones et al., 2017), sperm whale (Warren et al., 2017), and bottlenose dolphin (Neely et al., 2018) (Table 1). Scans for core genes from the BUSCO and CEGMA databases identified a near completeness of these genes in the assembly and support that we have largely reconstructed the entire genome. Only 122 scaffolds are needed, including the 34 largest scaffolds representing 50% of the whole genome, to almost completely cover the chromosomes of the *B. taurus* genome (Figure 1). The Circos plot depicting this (Figure 1) illustrates the near-completeness of the porpoise genome reached by theses 122 longest scaffolds and shows large regions of colinearity with the chromosome-level *B. taurus* genome assembly. Of these largest scaffolds some likely represent entire chromosomes, indicated by single *B. taurus* chromosomes, which are largely matched by only one or two porpoise scaffolds (e.g., cow chromosome 25, see Supplementary Information 1). Other *B. taurus* chromosomes are in only 2-3 pieces among our scaffolds, e.g. cow chromosomes 12 and 24 (Supplementary Information 1). Some of these fragmented matches likely indicate chromosomal rearrangements between domestic cattle and the harbour porpoise, two species diverged approximately 60Myrs ago within the Cetartiodactyla (Gatesy et al., 2013). Chromosomal rearrangements are seen several times among distinct lineage of Cetartiodactyla (Avila et al., 2015; Kulemzina et al., 2009, 2011; Pauciullo et al., 2014), for example between camel, pig and domestic cattle (Balmus et al., 2007). Based on these comparisons, we infer that our assembly represents a nearly complete genome of *P. phocoena* and that our largest scaffolds are nearly-complete chromosomes.

The draft annotation using Maker2 resulted in a total number of 22,154 annotated genes. This number is comparable to other published cetacean genomes, including 21,459 genes predicted in the bottlenose dolphin (Lindblad-Toh et al., 2011), 20,605 genes predicted in the minke whale (Yim et al., 2013), and 22,711 predicted genes in the grey whale (DeWoody et al., 2017). The annotated genes appear to broadly span key functional gene categories (biological processes, cellular components and molecular function), both across the annotated GO terms and the InterProScan results. Nonetheless this draft annotation should be further improved using transcriptome data (Fitak et al., 2016; Reyes-Chin-Wo et al., 2017) which were not available for our study organism. So far, with this draft annotation it is now possible to directly locate and further investigate key genes that may be under selection or underlie specific adaptive traits, both in cetaceans in general and in specific lineages (Chikina et al., 2016; McGowen et al., 2014; Yim et al., 2013). This genomic resource also provides candidate genes potentially underlying the local adaptation previously postulated within the harbor porpoise (Fontaine et al., 2014, 2017; Galatius et al., 2012). This includes genes with functions associated with low salinity and other cetacean specific traits (e.g., osmoregulation (aquaporines), blubber (ELOVL), reduction of smell, and taste receptors (Kishida et al., 2015; Pedro et al., 2015; Wang et al., 2015; Xue et al., 2014)). Sequence data available for gene families of interest will allow for comparative analyses across the cetacean/odontocete lineages, with a focus on porpoises representing the smallest cetaceans.

This genomic resource will also enable future genome-level analyses in the Phocoenidae, especially work that uses the *P. phocoena* genome to aid in genome assembly and annotation in other Phocoenidae species. This assembly contributes to the growing lists of whale genomes, and is the first sequenced genome for this family, which exhibits different ecological and morphological adaptations to the marine environment (Racicot et al., 2016; Wisniewska et al., 2016).

Our *P. phocoena* genome assembly will facilitate population genomic analyses within this species. It enables whole genome resequencing of further individuals to improve or revise subspecies and population delimitation (Rosel et al., 1999; Fontaine et al., 2014) or to allow identification of populations threatened by low effective population size/genetic isolation (Gui et al., 2013; Nadachowska-Brzyska et al., 2016; Park et al., 2015; Shafer et al., 2017). For the harbour porpoise this genome will further enable mapping of so far anonymous nuclear microsatellite (Wiemann et al. 2010) and SNP (Lah et al. 2016) loci, thus facilitating genomic monitoring of critically endangered population. It may be used for the compilation of informative SNP panels for population assignment in regions of population admixture, like in the Baltic region (Wiemann et al. 2010; Lah et al. 2016).

A first step in utilizing the *P. phocoena* genome was to investigate the demographic history of the harbor porpoise using the pairwise sequentially Markovian coalescent (PSMC) model. The PSMC estimates the demographic history of one diploid individual using the distance between locally estimated times to the most recent common ancestor (Li et al., 2011; Schraiber & Akey, 2015). Comparing the coalescence of alleles across the genome at a specific time points can indicate population contraction (many regions) or expansion (fewer regions). However this interpretation is only valid when one large panmictic population is assumed (Mazet et al., 2016), and it has been shown that population structure biases estimations of *N_e_* (Beichman et al., 2017; Li et al., 2011; Nadachowska-Brzyska et al., 2016). For example, the pattern of few large and many small pairwise differences could be observed both for populations that have experienced a bottlenecked, but also when a there is a multi-deem population structure (Nielsen & Beaumont, 2009). Following the recommendation of Li and Durbin (2011) to be cautious about inferences reaching further back in time than 1 Mio years, we concentrate here on the inference of potential population status between 20,000 and 1 Mio years ago (Nadachowska-Brzyska et al., 2016). Our sequenced specimen, sampled in the Baltic Sea, was assigned with high likelihood to the Beltsea population (Lah et al., 2016) by mtDNA analysis (exhibiting haplotype PHO 7; cf. Tiedemann et al., 1996; Wiemann et al., 2010). The Baltic Sea was formed only after the last glacial and – in its present form - is less than 10,000 years old (Andrén et al., 2011). Hence, for the time frame of our PSMC analysis, the sequenced specimen most likely represents the North Atlantic shelf population, a population now spanning from Iceland through the North Sea up to the Kattegat (Lah et al., 2016). Around 1 Mio years ago, our analysis infers an increase of *N_e_* from around 200,000 to an estimated 500,000 around 400,000 years ago, which could indicate an expansion of the Atlantic Shelf population, assuming a panmictic population. During the last interglacial period, the Eemian, the inferred *N_e_* remained relatively high. The abrupt inferred *N_e_* decline around 100kya ago, taken together with the formation of sea ice during the last glacial period could indicate a population bottleneck or a fragmentation into separate subpopulations (Nielsen et al., 2009). Among cetaceans (Brünchen-Olsen et al., n.d.; Foote et al., 2016; Yim et al., 2013; Zhou et al., 2013), the observed pattern, i.e., an increase in *N_e_* followed by a sudden drop, is most similar to the demographic history of bottlenose dolphin (*Tursiops truncatus*), a related species with a similar North Atlantic distribution. However, one should treat these inferences on historical *N_e_* with caution, as (1) they contain untested assumptions about panmixis, (2) other demographic scenarios could yield a similar pattern (Nielsen et al., 2009), and (3) the resolution of PSMC decreases for more recent times (Nadachowska-Brzyska et al, 2016).

In summary, we present here the first whole genome assembly and annotation of the harbour porpoise, at this point the most complete assembly for the Family *Phocoenidae*. This genome adds to available Cetacean genomes by supplying important resources for further investigation within the *Odontoceti* and *Cetacea*. This will provide an invaluable resource for further genetic studies of the harbour porpoise, both for whole-genome investigations into population structure and to identify key genes associated with local adaptation. This genome represents a crucial genetic resource for further investigation in the conservation, population genetics and phylogeny of other *Phocoenidae*, including the currently most rare marine mammal, the almost extinct Vaquita (*Phocoena sinus*) (Taylor et al., 2017).

## Acknowledgments

Financial support came from the Bundesamt für Naturschutz (FKZ # 3514824600), as part of a larger study of population genomics. We thank Prof. Dr. Michael Hofreiter for providing access to the Illumina NextSeq platform. Additional support came from the University of Potsdam. Large-scale computations were carried out on resources provided by Prof. Dr. Michael Lenhard at University of Potsdam and the High Performance Computing Cluster Orson2, managed by ZIM (Zentrum für Informationstechnologie und Medienmanagement) at the University of Potsdam.

## Data Accessibility

This Whole Genome Shotgun project has been deposited at DDBJ/ENA/GenBank under the accession PKGA00000000. The version described in this paper is version PKGA01000000. The draft annotation is deposited to Dryad (doi:10.5061/dryad.vr021gq).

## Authors Contributions

R.T. and L.L. designed the study; A.R. provided the sample and associated biological information. L.L. performed molecular lab work, S.H. performed initial *de novo* assembly, M.A. executed all genome annotations and analyses, M.A., S.H., A.B.D., and R.T. analyzed and interpreted the results, M.A. wrote the draft manuscript. All authors edited and approved the final manuscript.

---

**Supplement Material 1.** 122 *Phocoena phocoena* scaffolds mapped in the Circos plot, shown sorted to the *Bos taurus* chromosomes they match

| *Bos taurus*<br>chromosomes | *Number of*<br>*scaffolds*<br>*matched* | *Phocoena phocoena* scaffold IDs |
| --- | --- | --- |
| bt1 | 6 | Sc_11228, Sc_12227, Sc_1990, Sc_5358, Sc_9797, Sc_9900 |
| bt2 | 8 | Sc_11626, Sc_12227, Sc_13447, Sc_529, Sc_6515, Sc_6894, Sc_7661, Sc_9857 |
| bt3 | 9 | Sc_11018, Sc_1195, Sc_12865, Sc_3364, Sc_6515, Sc_7569, Sc_7648, Sc_7661, Sc_9681 |
| bt4 | 5 | Sc_12949, Sc_1454, Sc_2312, Sc_2531, Sc_6983 |
| bt5 | 6 | Sc_10530, Sc_10690, Sc_1183, Sc_4380, Sc_8015, Sc_8313 |
| bt6 | 7 | Sc_10623, Sc_10772, Sc_11919, Sc_12799, Sc_3795, Sc_4462, Sc_8081 |
| bt7 | 8 | Sc_1186, Sc_12016, Sc_12231, Sc_13164, Sc_2192, Sc_3821, Sc_7425, Sc_8211 |
| bt8 | 9 | Sc_11228, Sc_1608, Sc_3409, Sc_4380, Sc_4484, Sc_4874, Sc_515, Sc_664, Sc_880 |
| bt9 | 4 | Sc_10089, Sc_12949, Sc_9322, Sc_9860 |
| bt10 | 8 | Sc_10140, Sc_1186, Sc_12231, Sc_12865, Sc_2049, Sc_3337, Sc_7569, Sc_9860 |
| bt11 | 5 | Sc_10530, Sc_12437, Sc_12947, Sc_1608, Sc_7385 |
| bt12 | 2 | Sc_1065, Sc_11493 |
| bt13 | 6 | Sc_10356, Sc_12150, Sc_4275, Sc_8289, Sc_9915, Sc_9960 |
| bt14 | 5 | Sc_1753, Sc_3476, Sc_4138, Sc_7403, Sc_958 |
| bt15 | 5 | Sc_10137, Sc_10729, Sc_5614, Sc_6829, Sc_7410 |
| bt16 | 8 | Sc_11693, Sc_12029, Sc_141, Sc_2264, Sc_529, Sc_5715, Sc_8603, Sc_9900 |
| bt17 | 5 | Sc_10623, Sc_4462, Sc_5093, Sc_8289, Sc_8998 |
| bt18 | 4 | Sc_1099, Sc_3393, Sc_7023, Sc_9717 |
| bt19 | 4 | Sc_2056, Sc_7385, Sc_8323, Sc_9860 |
| bt20 | 3 | Sc_12231, Sc_7191, Sc_8211 |
| bt21 | 5 | Sc_12865, Sc_3249, Sc_3337, Sc_7569, Sc_9533 |
| bt22 | 5 | Sc_10627, Sc_12496, Sc_13313, Sc_4865, Sc_77 |
| bt23 | 2 | Sc_5169, Sc_8212 |
| bt24 | 2 | Sc_3090, Sc_3285 |
| bt25 | 1 | Sc_12150 |
| bt26 | 2 | Sc_7704, Sc_982 |
| bt27 | 2 | Sc_11228, Sc_11897 |
| bt28 | 3 | Sc_2616, Sc_9108, Sc_982 |
| bt29 | 5 | Sc_10137, Sc_10729, Sc_2203, Sc_5614, Sc_6829, |
| btx | 10 | Sc_10048, Sc_12502, Sc_2350, Sc_4277, Sc_5542, Sc_5764, Sc_6046, Sc_610, Sc_6806, Sc_933 |

